# Visible-Light-Controlled Residue-Selective Cross-Linking for Deciphering Protein Complexes and Dynamic Protein-Protein Interactions in Live Cells

**DOI:** 10.1101/2025.03.18.643847

**Authors:** Yali Xu, Hao Hu, Yu Ran, Wensi Zhao, An-Di Guo, Hui-Jun Nie, Linhui Zhai, Guang-Liang Yin, Jin-Tao Cheng, Shengna Tao, Bing Yang, Minjia Tan, Xiao-Hua Chen

## Abstract

<u>Cross-linking m</u>ass <u>s</u>pectrometry (XL-MS) has emerged as an attractive technology for investigating protein complexes and protein-protein interactions (PPIs). However, commonly used cross-linking strategies present significant challenges for precise analysis of protein complexes and dynamic PPIs in native biological environments. Here we present the <u>v</u>isible-light-controlled lysine-selective <u>cross-linking</u> (VL-XL) strategy for in-depth analysis of protein complexes and dynamic PPIs both *in vitro* and in live cells, building on light-induced primary amines and o-nitrobenzyl alcohols cyclization (PANAC) chemistry. We demonstrate that the VL-XL strategy effectively explores the dynamic dimerization of PD-L1 stimulated by exogenous modulators. Moreover, the VL-XL strategy successfully profiles the time-resolved EGF-stimulated EGFR interactome, providing valuable insights into the regulatory mechanisms of EGFR signaling and intracellular trafficking. Importantly, the VL-XL strategy efficiently deciphers the molecular glue (MG) induced dynamic PPIs and substrate profile of MG degrader, opening an innovative avenue for identifying neo-substrates. By harnessing the advantages of temporal controllability, good biocompatibility, and lysine selectivity, the VL-XL method simplifies MS data analysis and facilitates the acquisition of accurate structural information of protein complexes and the elucidation of elusive PPIs in live cells. Overall, the VL-XL strategy expands the XL-MS toolbox, and realizes in-depth analysis of protein complexes and dynamic PPIs, which will inspire innovative solutions for protein interactomes research and structural systems biology.

## Introduction

Elucidation of the protein complexes and dynamic protein-protein interactions (PPIs) is crucial for understanding of understudied proteins and molecular basis of cell functional diversity, as well as providing promising targets for drug discovery.^1-3^ Indeed, structure-focused techniques (e.g. NMR spectroscopy, cryo-electron microscopy) have provided stunning structural images of isolated proteins and their complexes.^1, 4, 5^ By contrast, deciphering protein conformational changes, protein networks and dynamic PPIs within cellular milieu remains an important yet largely unexplored object in the field of structural systems biology, protein interactomes and fundamental biology.^1, 4, 5^ To this end, cross-linking mass spectrometry (XL-MS) has been developed recently to probe protein conformations and dynamic PPIs.^1, 4, 5^ Compared with other strategies, XL-MS has unique ability to covalently lock interplayed proteins (even for weakly or transiently interacting partners) in native status and identify their physical contacting sites.^1, 2, 4-6^ Therefore, XL-MS has emerged as an attractive technology to provide new insights into conformation and dynamics of protein complexes^7, 8^ involved in various biological processes, as well as PPIs in organelles^9, 10^ and at whole-proteome scale^11-13^ in living cells.

In XL-MS technology, the chemistry of the cross-linking is crucial for acquiring accurate information of protein complexes, as well as the depth of protein interactions and dynamics.^1, 2, 4, 5^ With regard to the chemistry of cross-linking, there are two kinds of cross-linking strategies, spontaneous reactive functionalities and photo-controlled functionalities.^1, 4, 5^ Remarkably, spontaneous reactive cross-linkers based on lysine-targeting N-hydroxysuccinimide (NHS) esters (Figure 1a) are the most widely used due to the high abundance of lysine residues on protein surface.^1, 4, 5^ To date, large amount of homobifunctional NHS-ester-related cross-linkers, and heterobifunctional cross-linkers integrating NHS-ester moiety with photo-inducible moiety moiety (Figure 1a) have been developed, and successfully employed for understanding structures of protein assemblies^7, 8^ and network of intracellular PPIs.^9-13^ However, NHS-ester-related cross-linkers have several inherent limitations (Figure 1a). The NHS-ester moieties are susceptible to hydrolysis compromising the cross-linking efficiency and limiting the applications in living cells.^14, 15^ In addition, unexpected side reactions with hydroxyl groups on Ser/Thr/Tyr (∼ 30% frequency) would significantly complicate the data interpretation for proteome-wide cross-linking MS analysis.^5, 6, 15^ On the other hand, the incorporated conventional photo-cross-linking functionalities (such as diazirines, azides, and benzophenones) in heterobifunctional NHS-ester-related cross-linkers target nearly all residues non-selectively when activated by ultraviolet (UV) light (Figure 1a, lower panel), generating excessive and high-density cross-links, significantly complicating the MS analysis and artificially distort protein structural information.^4, 5^ Furthermore, these abovementioned NHS-ester-related cross-linkers are spontaneous reactivity lacking temporal controllability,^14, 15^ thus, hamper the in-depth exploration of protein complexes, interactomes and dynamics in various biological processes, in living systems.

**Figure 1.**
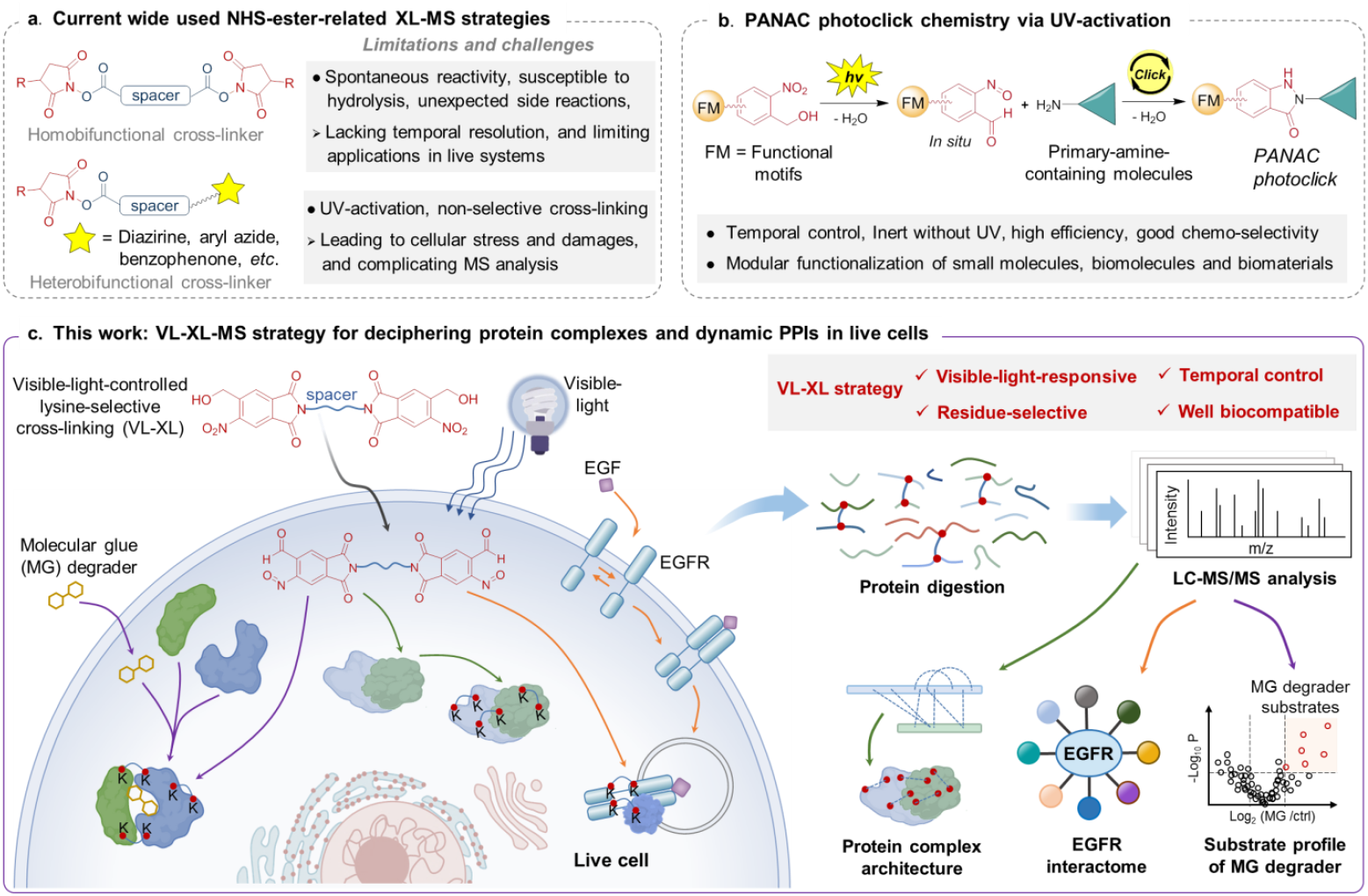
Schematic representation of currently used cross-linkers, and design of VL-XL strategy for analysis of protein complexes and PPIs. **a**) Currently widely used NHS ester-related XL-MS strategies. **b**) The multiple characteristics and applications of PANAC photoclick chemistry. **c**) Design of VL-XL-MS strategy to cross-link protein complexes, and decipher protein complex architectures and dynamic PPIs *in vitro* and in live cells.

Indeed, the light-activation is an ideal way to realize temporal and spatial control in biological processes.^16^ In contrast to homobifunctional NHS-ester-related cross-linkers, the photo-cross-linkers tethering two photo-reactive moieties remain largely underdeveloped.^4, 5^ A major challenge in development of the bifunctional photo-cross-linkers for XL-MS is the non-selective cross-linking of any nearby residue of protein complex from two photo-reactive functionalities,^17, 18^ which will dramatically complicate database searches, causing more false positives and reducing peptide identification sensitivity.^4^ Until recently, only photocaged quinone methide-based cross-linker was developed by Wang for XL-MS studies, in which the homobifunctional photo-responsive functionalities target multiple residues, induced by UV light-activation.^19^ Nevertheless, all cross-linkers derived from photo-reactive functionalities (Figure 1a, lower panel and quinone methide-based cross-linker) for XL-MS studies so far are activated by UV light.^1, 4, 5, 19^ In fact, UV irradiation can cause unwanted photochemical transformations and activate DNA damage response pathways,^20^ such as oxidation of multiple residues in proteins,^21^ and the ribotoxic stress response,^22^ leading to significant cellular stress and damages, thus, hindering the precise characterization of protein interactions and applications in live systems.^21-23^ Therefore, it remains a challenge and an unmet need in XL-MS field to develop efficient and well biocompatible cross-linking strategy, to achieve high accessibility for precise MS data analysis and in-depth protein complexes investigation, as well as temporal controllability for dynamic PPIs analysis in live systems.

Herein, we have developed a <u>V</u>isible-light-controlled <u>L</u>ysine-selective <u>Cross-Linking</u> (VL-XL) strategy to cross-link protein complexes, inspired by our previous PANAC photoclick chemistry (Figure 1b),^24-27^ enabling in-depth analysis of protein complexes and dynamic PPIs both *in vitro* and in live cells (Figure 1c). We demonstrated that the VL-XL method effectively explores dynamic PD-L1 dimerization stimulated by an exogenous modulator, and profiles time-resolved EGF-stimulated EGFR interactome providing valuable insights into the regulatory mechanisms of EGFR signaling and intracellular trafficking. Moreover, the VL-XL strategy is capable of deciphering substrate profile of E3 ligase cereblon (CRBN) induced by molecular glue (MG), leading to the discovery of the neo-substrates, opening an innovative avenue for identifying neo-substrates of MG degraders. Thus, by harnessing the advantages of temporal controllability, lysine selectivity and good biocompatibility, the VL-XL method provides residue-defined cross-linked sites with high confidence and simplifies MS data analysis, facilitating acquiring accurate information of protein complexes and in-depth investigation of elusive dynamic PPIs in native biological environments. Overall, the VL-XL strategy expands the XL-MS toolbox and offers a reliable cross-linking method for XL-MS and protein interactomes research.

## Results and Discussion

### Design of light-responsive and lysine-selective photo-cross-linkers for XL-MS

Recently, we have developed a photoclick chemistry, light-induced <u>p</u>rimary <u>a</u>mines and o-<u>n</u>itrobenzyl <u>a</u>lcohols <u>c</u>yclization (PANAC, Figure 1b).^24-27^ The o-nitrobenzyl alcohol (o-NBA) derived reactants are inert before light activation, and the o-NBA moiety can be activated by UV light into reactive o-nitrosobenzaldehyde intermediate which is selectively conjugated with primary amines. The PANAC photoclick chemistry features fast kinetics, high chemo-selectivity^24, 25, 28^ and temporally-controlled manner (Figure 1b). Practically, the light-induced PANAC reaction is highly efficient with low concentrations of reactants under operationally simple and mild conditions, without the use of toxic metal catalysts and ligands *in vitro* and in living systems, conferring less toxicity to biological systems.^24^ Harnessing these advantages, we and other groups have applied the PANAC photoclick chemistry for modular functionalization of diverse small molecules,^29-31^ biomolecules^24-28, 32-34^ and biomaterials.^35-37^

Based on these insights into PANAC photoclick chemistry and inspired by the success of visible-light-powered chemical transformations, we recently questioned whether it might be possible to merge the advantages of PANAC photoclick chemistry with the good biocompatibility of visible light,^23, 38^ to develop a visible-light-controlled lysine-selective cross-linking (VL-XL) strategy for XL-MS analysis (Figure 1c), to overcome long-standing challenges associated with spontaneous reactive cross-linking and conventional UV-driven non-selective photo-cross-linking (Figure 1a). We have designed a lysine-selective photo-cross-linker incorporating the light-inducible o-NBA moiety with appropriate linker compositions (Supplementary Figure 1) to facilitate the temporally controlled cross-linking of protein complexes and protein–protein interactions (PPIs). The UV-light-induced photo-cross-linking strategy has been successfully employed to investigate protein complexes and PPIs both in vitro and in living cells. We envisioned that further structure design and modification of the o-NBA moiety would realize the development of visible-light-responsive photo-cross-linkers for XL-MS studies (Figure 1c). Consequently, the VL-XL strategy would confer the residue-selective cross-linking and superior biocompatibility, thus, simplifies the MS data analysis and acquiring accurate information of protein complexes, as well as conferring temporally-controlled manner for in-depth analysis of protein complex or dynamic PPIs in living systems.

### Development of visible-light-controlled cross-linkers for XL-MS

By structural modification and optimization of the o-NBA moiety, we found that the introduction of a carbonyl group bridging the phenyl group and amide nitrogen extends the absorption wavelength to visible-light range (Supplementary Figure 9). To our delight, the model compound 5-(<u>h</u>ydroxymethyl)-2-<u>m</u>ethyl-6-<u>n</u>itroisoindoline-1,3 -<u>d</u>ione (HMND) rapidly reacted with lysine derivative (Cbz-lys-OMe), generating the expected indazolinone product with high conversion (> 95%) under visible-light activation (420-430 nm, Supplementary Figure 10). With above obtained cross-linking profiles and spacer diversity of the cross-linkers in hand, we designed and developed two visible-light (VL)-inducible cross-linkers (Figure 2a, namely VL-CO cross-linker or VL-CO2 cross-linker, respectively).

**Figure 2.**
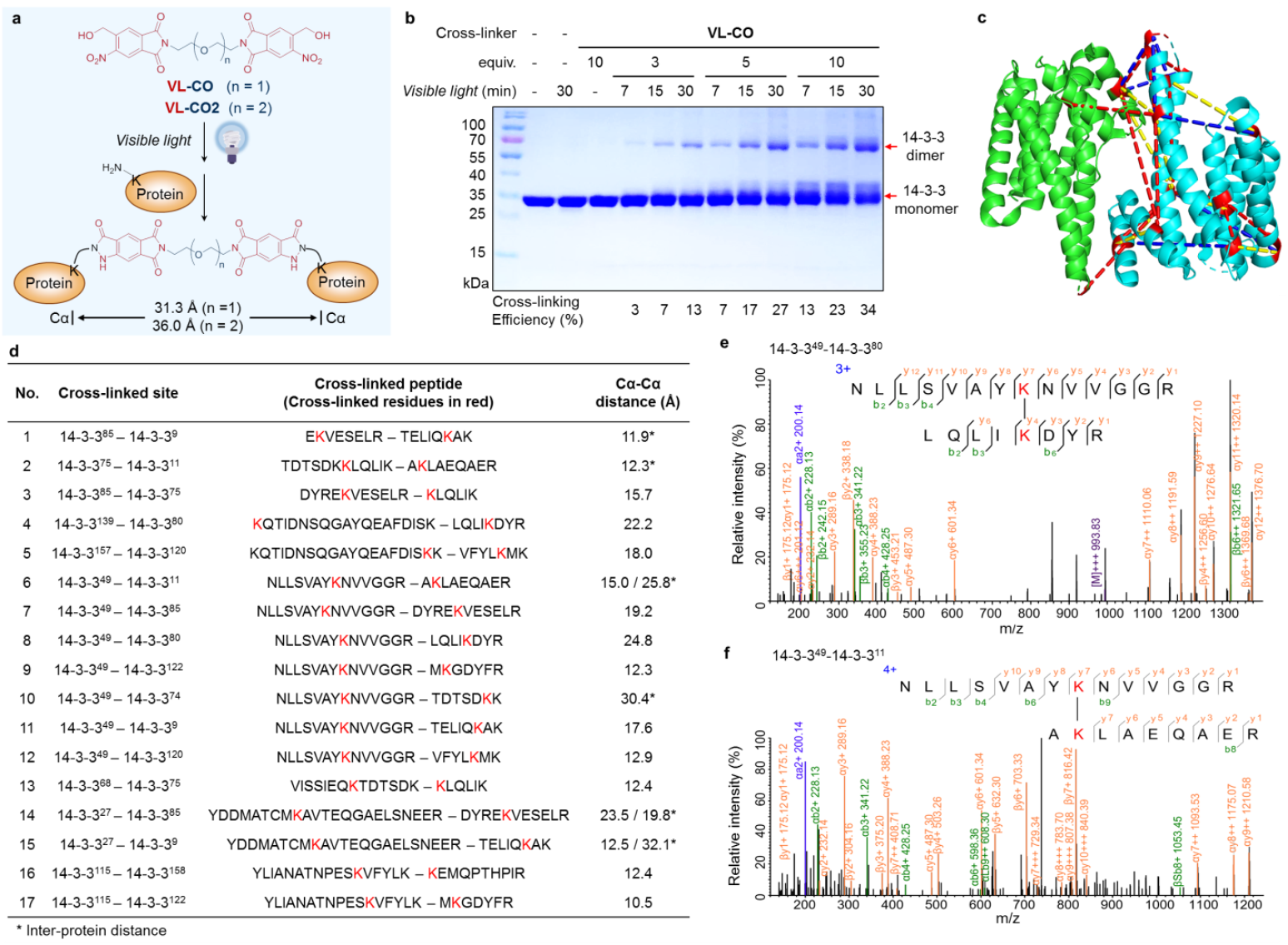
VL-XL cross-linker-mediated cross-linking on purified 14-3-3 protein. (a) Scheme of VL-CO and VL-CO2 structures and visible-light-induced cross-linked products. (b) SDS-PAGE analysis of dimeric cross-linking of 14-3-3 (10 μM) with indicated concentration and exposure time of visible light. (c) Comparison of sites cross-linked by VL-CO and VL-CO2, mapped onto the crystal structure of 14-3-3 protein (PDB ID: 2BTP). The Cα-Cα uniquely cross-linked by VL-CO are connected with red dashes; uniquely cross-linked by VL-CO2 with blue dashes; cross-linking sites by both of these cross-linkers with yellow dashes. For the ease of viewing, the intra-protein cross-links are mapped onto one monomer (cyan). (d) High-confidence cross-linked peptides and sites identified from VL-CO cross-linking. For Cα–Cα distance, * indicates inter-protein distance, otherwise, intra-protein distance. (e) Representative tandem mass spectrum of intra-protein cross-link identified from VL-CO cross-linking. (f) Representative tandem mass spectrum for inter-protein cross-link identified from VL-CO cross-linking.

To our delight, VL-CO successfully cross-linked the 14-3-3 protein under visible-light activation (420-430 nm). From the SDS-PAGE analysis (Figure 2b), this visible-light-responsive cross-linker provides good cross-linking efficiency (more than 30%), and the visible-light-induced cross-linking yield increased along with the cross-linker concentration and irradiation time (Figure 2b). Then, the cross-linked protein complexes were subjected to tandem MS analysis. By manual inspection of CSMs, we identified 17 pairs of high-confidence cross-linked sites (Figure 2c,d and Supplementary Figure 11). All the Cα–Cα distances of these cross-linked sites complied with the distance constraint (31.3 Å, Figure 2a,c,d), and indicated good structural compatibility of the identified cross-linking sites via cross-linker VL-CO, suggesting the potential of our cross-linker for accurate structural studies of the proteins of interest.

Next, we further performed cross-linking experiments and analysis on VL-CO2 (Figure 2a), which has longer and more flexible spacer compared to VL-CO. As expected, the cross-linking yield of VL-CO2 increased along with the cross-linker concentration and irradiation time (Supplementary Figure 12a), and was comparable to VL-CO (Figure 2b). From MS analysis, we identified 14 pairs of high-confidence cross-linked sites (Supplementary Figure 12b,13) and all the Cα– Cα distances are smaller than the distance constraint (36.0 Å, Figure 2a), suggesting these cross-linked sites of VL-CO2 cross-linking were highly compatible with the crystal structure of 14-3-3 protein. Interestingly, among them, 7 cross-links (Figure 2c, yellow dashes) were the same as those captured by the cross-linker VL-CO (shorter spacer cross-linker), indicating different cross-linked-site coverage of two VL-XL cross-linkers. Altogether, these results demonstrated that the cross-linking mediated by the VL-XL cross-linkers was successfully activated by visible-light with temporal controllability, and the structure variation of VL-XL cross-linkers could provide richer information for accurate analysis of PPIs and conformations of protein complexes, in certain research scenarios.

### VL-XL strategy for profiling of EGF-stimulated EGFR interactome by label-free quantitative proteomics

The CLogP values of VL-XL cross-linkers are within the range of those of reported cross-linkers that are widely used for in-cell cross-linking, suggesting their cell permeability (Supplementary Figure 14). In addition, preliminary experiments showed that short-term incubation of unactivated VL-XL cross-linkers or visible-light irradiation had no apparent effect on cell viability (Supplementary Figure 15). These results indicated the VL-XL strategy is biocompatible and suitable for studying cellular PPIs. In this context, we first explored the applicability of VL-XL strategy with visible-light activation in live cells. We applied our VL-XL method for cross-linking of 14-3-3 dimer protein (Supplementary Figure 16a) and the PD-L1 dimerization induced by BMS1166 (Supplementary Figure 16b,c) in 293T cells. Based on the analysis of these results, to our delight, we successfully demonstrated that the VL-XL strategy has the ability to cross-link protein complex and investigate dynamic PPIs in response to exogenous stimulus in live mammalian cells. Encouraged by these results, we reasoned that the VL-XL strategy is capable of profiling PPIs via covalent cross-linking to effectively capture weak or transient interactions which are often lost in conventional co-immunoprecipitation (co-IP). Thus, the VL-XL strategy would offer a higher coverage of PPI network in live cells.

Then, as a proof-of-concept study, we implemented VL-XL-immunoprecipitation (VL-XL-IP) to capture proteins that bind with EGFR following EGF stimulation, and benchmarked this method with co-IP (Figure 3a). As the EGFR interactome is well characterized, validation of the pulldown data from VL-XL-IP or co-IP can be readily carried out by cross-referencing with the inventory of known EGFR interactors.

**Figure 3.**
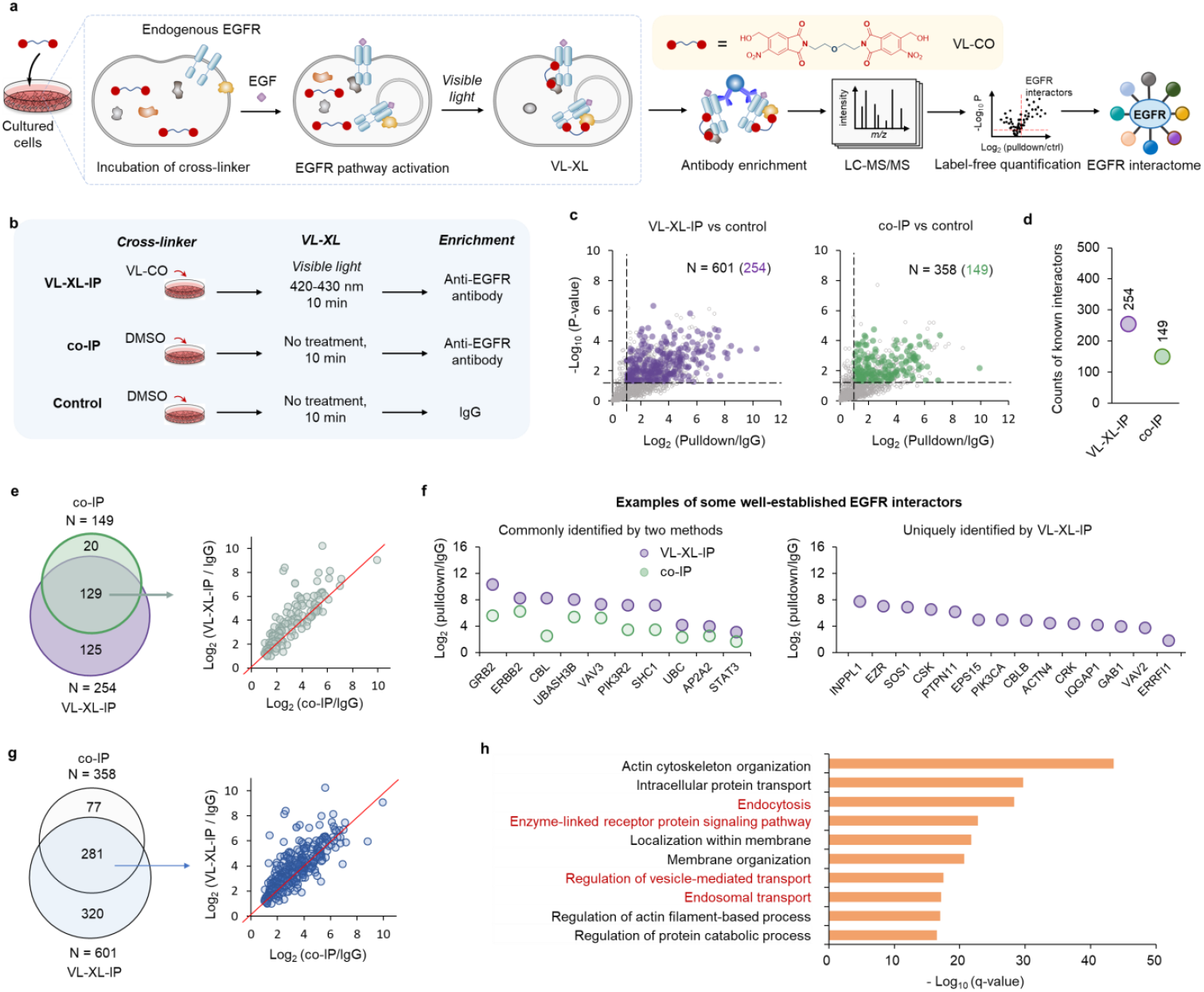
Comparing the performance of VL-XL-IP and co-IP in profiling the EGFR interactome by label-free quantitative proteomics. **a**) Following incubation with VL-CO for 30 min and with EGF for 5 min, HeLa cells were subjected to light-activation (420-430 nm). After cell lysis, EGFR-interacting proteins were enriched by anti-EGFR IP, and identified by label-free quantitative proteomic analysis. **b**) Design of VL-XL-IP, co-IP and control experiments. **c**) Scatter plots showing the proteins significantly enriched by VL-XL-IP or co-IP compared to IgG. Significant enrichment: pulldown/IgG intensity ratio > 2 and p < 0.05 (n = 3). Reported EGFR interactors were colored purple (in VL-XL-IP) or green (in co-IP). **d**) The number of reported EGFR interactors enriched. **e**) Left panel: comparing the coverage on reported EGFR interactors enriched by VL-XL-IP and co-IP. Right panel: comparing the enrichment ratios of reported EGFR interactors commonly enriched by two methods. **f**) Enrichment ratios of some well-established EGFR interactors. **g**) Comparison of all proteins enriched by VL-XL-IP and co-IP, illustrated by a Venn diagram (left) showing coverage and a scatter plot (right) comparing enrichment ratios of overlapped proteins. **h**) Top 10 significantly enriched GO-BP terms for proteins enriched by VL-XL-IP. Terms for well-characterized EGFR-involved biological processes are colored red.

In VL-XL-IP experiment, we first treated the serum-starved cultured cells *in situ* with the newly developed VL-CO cross-linker for 30 min to achieve equilibrium (Figure 3b). Notably, the photo-cross-linkers remain inactive during incubation, making them compatible with the original culture media (containing millimolar-level concentration of amino acids), without need to switch to an amino-free medium (such as PBS) as often required for NHS-based cross-linkers.^11-13^ Since the incubation occurs in the same environment where the cells are grown, we reason that it preserves the native and physiological cellular interactions. Following the incubation with VL-CO cross-linker, cells were subjected to 5-min stimulation with EGF and then exposed to visible light for cross-linking (Figure 3b). The co-IP experiment was conducted using the same procedures, but without the addition of photo-cross-linker and light-activation (Figure 3b). In addition, we set a control group in which IgG-immobilized beads were used to incubate with cell lysate for quantitative comparison to eliminate false-positive results arising from non-specific binding to the antibody or the beads (Figure 3b).

We first evaluated the reported EGFR-interacting proteins enriched by VL-XL-IP and co-IP. A remarkably higher number proteins are observed in VL-XL-IP (254) compared to co-IP (149) (Figure 3c,d and Supplementary table 1,2). In regard to the coverage, VL-XL-IP not only covers approximately 87% of the proteins enriched by co-IP (129 out of 149), but also expands the coverage by around 84% (125 extra proteins) compared to that of co-IP (149 proteins) (Figure 3e, left panel). For the 129 proteins in the overlap, most proteins exhibit higher enrichment ratios in VL-XL-IP compared to those in co-IP (Figure 3e, right panel), such as well-characterized EGFR signaling pathway-related interactors GRB2, ERBB2, PIK3R2, SHC1, STAT3, as well as endocytosis and degradation-related interactors CBL, UBASH3B, VAV3, UBC, AP2A2, etc (Figure 3f, left panel). Among the 125 EGFR interactors uniquely enriched by VL-XL-IP, we also observed a number of well-characterized EGFR interactors with high enrichment ratios, such as INPPL1, EZR, SOS1, CSK, PTPN11, EPS15, PIK3CA, CBLB, ACTN4, CRK, IQGAP1, GAB1, VAV2, ERRFI1, etc (Figure 3f, right panel).

With regards to all proteins enriched by two methods, we also found a larger number of proteins (601) were enriched by VL-XL-IP, which covers around 78% (281) of the proteins enriched by co-IP (358) (Figure 3g). Generally speaking, for a given protein that is commonly enriched by two methods, the enrichment ratio observed in VL-XL-IP is typically higher than that seen in co-IP (Figure 3g). Gene Ontology – biological processes (GO-BP) enrichment analysis shows that proteins enriched by VL-XL-IP are indeed involved in well-characterized EGFR-related pathways, including endocytosis, enzyme-linked receptor protein signaling pathway, regulation of vesicle-mediated transport, endosomal transport, etc (Figure 3h). Collectively, these findings highlight the enhanced sensitivity and coverage of the VL-XL strategy in the profiling of the EGF-stimulated EGFR interactome.

### Time-resolved and proteome-wide analysis of EGF-stimulated dynamic EGFR interactome by VL-XL strategy coupled with multiplexing quantitative proteomics

EGF binding promotes EGFR dimerization and autophosphorylation, which activates various downstream signaling pathways. In the meantime, the activated EGFR undergoes internalization and endocytic trafficking, ultimately being degraded in the lysosome or recycled back to the plasma membrane.^39^ Although these processes are extensively studied, the EGF-stimulated PPIs of EGFR are rarely investigated in a proteome-wide and time-resolved manner, thus the related regulatory mechanism remains not fully understood. The light-controllability of the VL-XL allows for the precise capture of transient or time-specific PPIs that may change rapidly over time, thus providing an avenue for acquiring critical insights into the binding kinetics of the EGFR interactome. Therefore, we then employed VL-XL strategy for time-resolved analysis of the EGF-stimulated dynamic interactome of EGFR.

Cells were incubated with VL-CO to achieve equilibrium, treated with or without EGF for varied time courses (2, 10, 20, 60 min), and subjected to visible-light activation for cross-linking. After cell harvest and lysis, EGFR interactors were enriched, tryptically digested, tandem mass tag (TMT)-labeled for LC-MS/MS analysis (Figure 4a). We found that 115, 52, 62 and 95 proteins exhibited significantly upregulated interactions with EGFR at 2, 10, 20, 60 min after EGF stimulation, respectively (Figure 4b and Supplementary Figure 17a). This corresponds to a total of 152 EGFR-interacting proteins (Supplementary table 3), half of which were previously unreported. We first analyzed the kinetics of several well-established EGFR interactors including the EGFR signaling pathway-related SHC1, SOS1, GRB2, PIK3CB, PIK3R2 and EGFR endocytosis-related AP2M1 (Figure 4c and Supplementary Figure 17b). We found that their kinetics are consistent with the results from a previous study,^40^ corroborating the effectiveness of our methods.

**Figure 4.**
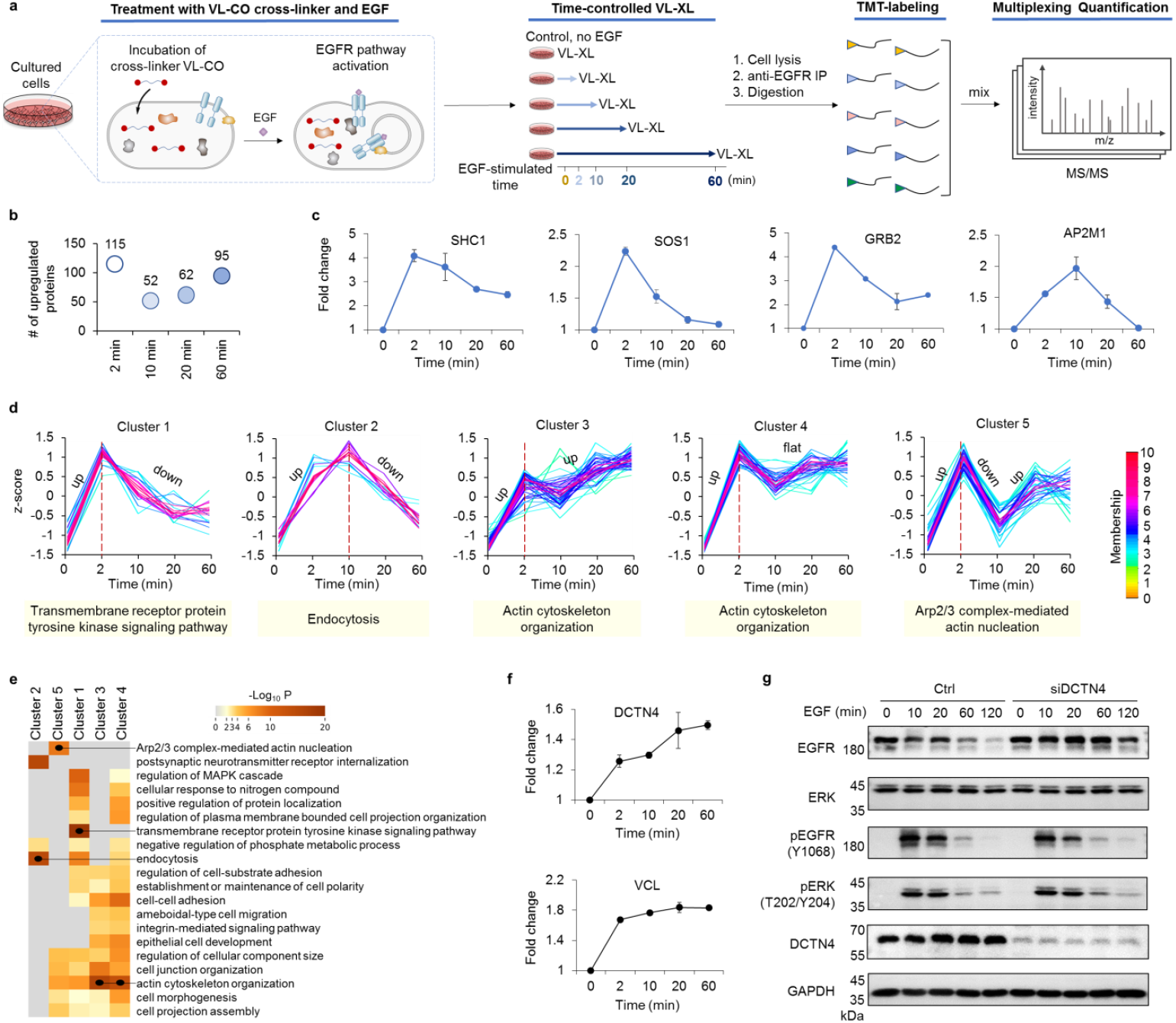
Application of VL-XL coupled with TMT-based multiplexing quantitative proteomics in time-resolved analysis of EGF-stimulated dynamic interactome of EGFR. **a**) HeLa cells pre-incubated with VL-CO were treated with or without EGF (control), followed by light irradiation.After cell lysis, EGFR-interacting proteins were enriched by anti-EGFR IP, digested, labeled with distinct channels of TMT, analyzed by MS-based multiplexing quantification. **b**) The number of proteins significantly upregulated (fold change > 1.2 and p < 0.05) at post-EGF time points compared to the control (0 min) from biological duplicates. **c**) Binding kinetic curves of representative known EGFR interactors in response to EGF stimulation. Error bars represent standard deviations (n = 2). **d**) Clustering of the interacting kinetic curves of proteins that were significantly upregulated in at least one time point. Fold change values were z-score transformed and classified into 5 clusters using fuzzy c-means algorithm. Below each cluster, the most significantly enriched GO-BP term is displayed. **e**) Heatmap showing GO-BP enrichment analysis for proteins in five clusters, with the most significantly enriched GO term in each cluster marked by a black dot. **f**) Kinetic curves of DCTN4 or VCL binding with EGFR after EGF stimulation. Error bars represent standard deviations (n = 2). **g**) Western blot analysis of protein level of EGFR, ERK, DCTN4, VCL, and phosphorylation of EGFR (pY1068) and ERK (pT202/pY204) in siDCTN4-knockdown group and control (ctrl) group.

Then, we set out to characterize the overall binding kinetics of the 152 EGF-upregulated proteins. The z-score-normalized binding kinetic curves of 152 proteins were classified into five clusters (Figure 4d and Supplementary table 4), and a GO-BP enrichment analysis was performed for each cluster (Figure 4e). In cluster 1, proteins reach their peaks at 2 min, and EGFR-related signaling pathway (e.g. transmembrane receptor protein tyrosine kinase signaling pathway) are most significantly enriched, including ERBB2, GRB2, PIK3CA, PIK3CB, PIK3R2, CRK, CRKL, PTPN11, SHC1, SOS1, etc. In cluster 2 where curves peak at 10 min, the most significantly enriched terms are endocytosis and receptor internalization, mainly involving the AP2 complex components AP2A1, AP2A2, AP2B1 and AP2M1. For these two clusters, the binding kinetics and enriched GO-BP terms are highly reflective of the established EGFR-related biological processes upon EGF stimulation. In cluster 3 and 4, proteins exhibit rapid association with EGFR at the beginning, and then slowly increase (cluster 3) or reach a plateau (cluster 4) at longer time course. For these proteins, some actin-related processes (e.g. actin skeleton organization, cell adhesion, cell junction organization, etc.) are significantly enriched, which is consistent with the critical roles of actin skeleton in intracellular protein transport. In cluster 5, proteins display multiple kinetic phases (i.e. rapidly binding to EGFR within the first 2 min, dissociating from 2 and 10 min, and then gradually re-associating over a longer time course), and the most significantly enriched GO-BP term is Arp2/3 complex-mediated actin nucleation (including ARPC4, ARPC1B, ACTR2, ARPC5L). Since the Arp2/3 complex plays a crucial role in initiating the synthesis of branched actin,^41^ the multi-phase kinetic curves suggest the formation of actin network is involved in both early endocytosis and late-phase events of EGFR intracellular trafficking. Altogether, the time-resolved analysis not only identified new EGFR interactors with determined binding kinetics, but also revealed various binding kinetic trends and their associated biological processes invoked upon EGF stimulation. These results will provide valuable insights into the regulatory mechanisms of EGFR signaling and intracellular trafficking.

### Discovery of the DCTN4 and VCL affecting EGF-stimulated degradation of EGFR

Interestingly, the EGF-stimulated EGFR interactome comprises many proteins associated with actin cytoskeleton organization process (Figure 4d, cluster 3 and 4). For example, we found two proteins, DCTN4 (also known as p62) and vinculin (VCL), interacted with EGFR during the whole investigated time course (Figure 4f). DCTN4 is an essential subunit of the dynactin which drives the cellular transport of a wide variety of cargoes, including endosome and lysosome.^42^ VCL can associate with the early endosome marker Rab5,^43^ suggesting a role in endocytosis. As demonstrated in *in sit*u cross-linking experiment, compared to IgG group, the VL-XL-IP group showed a protein band with molecular weight indicating cross-linked EGFR-VCL complex, validating their interaction in cells (Supplementary Figure 21a). Since DCTN4 and VCL have not been reported to interact with EGFR previously, their regulatory roles on EGFR remain unclear.

According to their known cellular functions related to endosome, we hypothesized that DCTN4 and VCL likely participated in the process from internalization to lysosome-degradation of EGFR. Thus, we knocked down DCTN4 and VCL in HeLa cells, and analyzed their effects on the EGF-induced temporal alteration of EGFR protein level and downstream phosphorylation by Western blot (Figure 4g and Supplementary Figure 17c). In control group (treated by negative control siRNA, siCtrl), protein levels of EGFR profoundly decreased under longer (60 min and 120 min) treatment of EGF. In siDCTN4- or siVCL-treated groups, prolonged treatment (60 and 120 min) of EGF did not substantially reduce the protein levels of EGFR compared to untreated sample (i.e. 0 min), indicating the knockdown of these two proteins substantially slowed the EGFR degradation. Whereas, the temporal pattern of the phosphorylation of EGFR (pY1068) and its downstream effector ERK1/2 (pT202/pY204) were similar between the knockdown (siDCTN4 or siVCL) group and control (siCtrl) group. These results together suggest that DCTN4 and VCL regulate the EGF-stimulated degradation of EGFR, but not the phosphorylation signaling pathway. Furthermore, based on identified cross-linked peptides and prediction of PPIs with AlphaFold 3 ^44^ (Supplementary Figure 19), we provide structural insights into the topology of EGFR/VCL and EGFR/ DCTN4 protein complexes.

### Deciphering the substrate profile of MG degrader in live cells via VL-XL strategy

The development of MG degraders holds great significance as it offers a novel approach to target previously “undruggable” proteins, expanding therapeutic possibilities beyond traditional inhibitors.^45, 46^ Despite many strategies have been developed, it remains a substantial challenge to identify the elusive substrates of MG degraders.^47, 48^ As degrader-induced interaction between the substrates and E3 ligase is the initial step of targeted protein degradation, profiling these interactions provides an avenue for identifying new degradation substrates. Given that such interactions can be weak and transient, we reasoned that our VL-XL method, which enables irreversibly locking of cellular interactions *in situ*, offers an effective way to capture them. To demonstrate the ability of VL-XL strategy for profiling MG-induced substrates, we set out to apply our method to investigate the repertoire of endogenous degradation substrate of compound CC-885, a well-known MG that induces the degradation of GSPT1 in an E3 ligase cereblon (CRBN)-dependent manner.^49^

Human 293T cells overexpressing CRBN with Twin-Strep tag were incubated with VL-CO cross-linker and CC-885, followed by visible-light activation. After cell lysis, proteins were affinity-enriched by Strep-Tactin beads, tryptically digested, and analyzed by LC-MS/MS (Figure 5a). CC-885-induced CRBN-interacting proteins were identified by comparing the CC-885-treated group to the DMSO-treated group (control) with label-free quantification (cutoff: CC-885/control enrichment ratio > 2, p < 0.05, Supplementary table 5, 6). Notably, the known substrates GSPT1 (Supplementary Figure 18) and its close homolog GSPT2 exhibited largest ratios (Figure 5b). *In vitro* XL-MS experiments identified cross-linked peptides of CRBN and GSPT1 (Supplementary Figure 20), supporting our VL-XL strategy could provide direct evidence of PPIs. We also identified VCL, which is recently reported as a new but non-degradable substrate of CRBN induced by CC-885,^50^ and found it to be significantly enriched (Figure 5b). Western blot analysis provides evidence for *in situ* cross-linking of CRBN-GSPT1 and CRBN-VCL complexes in cells, induced by MG degrader (Supplementary Figure 21b). These results support the specificity and effectiveness of our method. Furthermore, we confirmed that SESN2, a highly conserved stress-inducible metabolic protein that influences cancer development in a multifaceted way,^51^ exhibited time-dependent degradation upon CC-885 treatment (Figure 5b, c, d). The degradation was rescued by MLN7243, MLN4924 and MG132 (inhibiting the activities of E1 ubiquitin-activating enzyme, CUL4-based E3 ligase, and proteasome, respectively), suggesting the degradation was dependent on the ubiquitin-CUL4 E3 ligase-proteasome pathway (Figure 5e). SESN2 was also degraded in triple-negative breast cancer MDA-MB-231 and MDA-MB-468 cells upon the treatment of CC-885 (Figure 5f), which indicates SESN2 is a neo-substrate of the MG degrader CC-885. Taken together, these results demonstrated that the protein SESN2 could be targeted and regulated by MG degrader through degradation in cancer cells.

**Figure 5.**
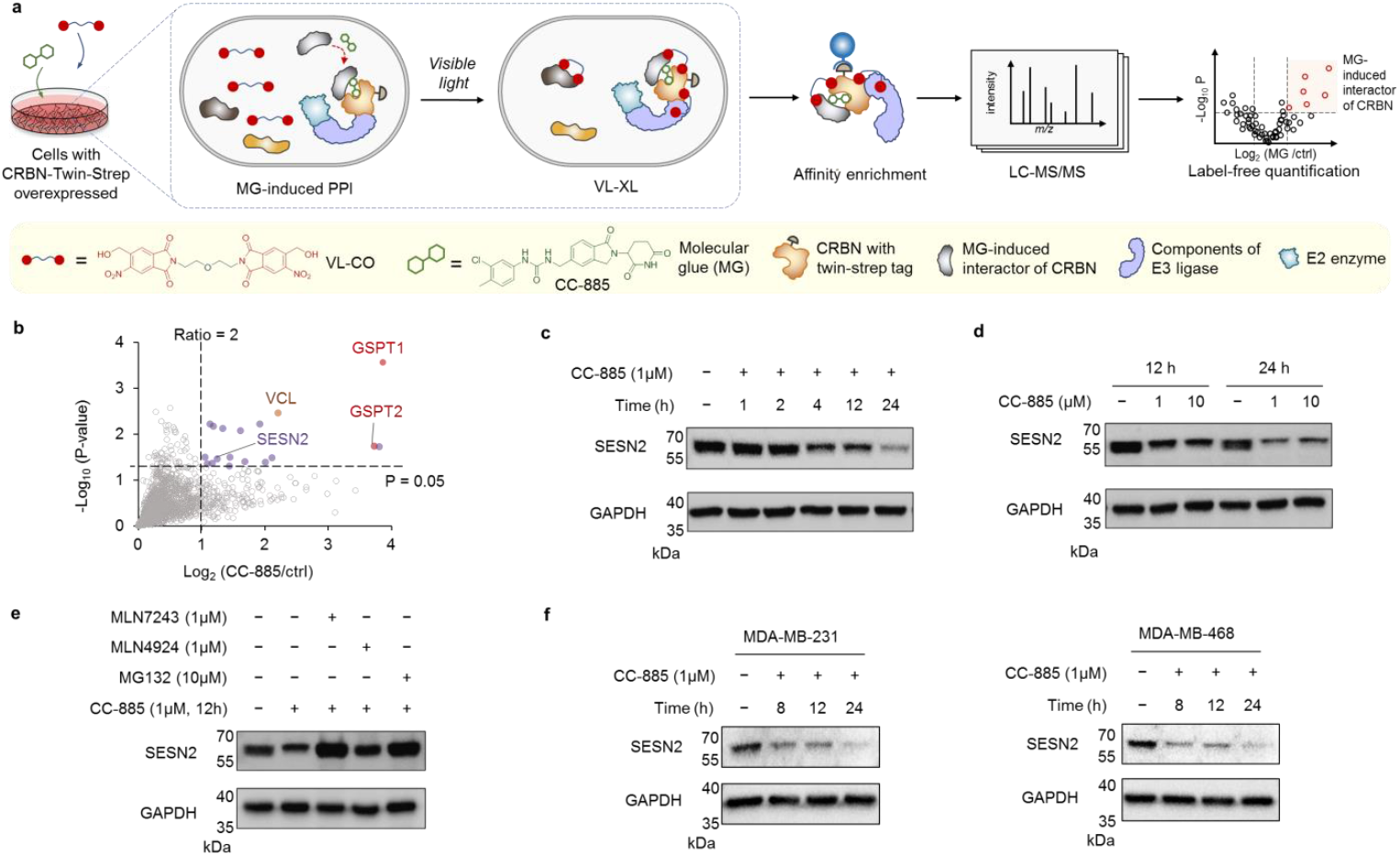
Application of VL-CO-based VL-XL for identifying the neo-substrate of molecular glue CC-885. **a**) 293T cells preincubated with VL-CO and CC-885 were subjected to light irradiation (420-430 nm). After cell lysis, interactors of CRBN were enriched and analyzed by label-free quantitative proteomics. **b**) Proteins significantly enriched in CC-885-treated group compared to control (ctrl) group. Significant enrichment: CC-885/Ctrl intensity ratio > 2 and p < 0.05 (n = 3). **c**) Western blot analysis of time-dependent degradation of SESN2 in 293T cells. **d**) Western blot analysis of the degradation of SESN2 at two concentrations of CC-885 over two different time periods in 293T cells. **e**) Western blot analysis of the effect of MLN7243, MLN4924, MG132 on the CC-885-induced degradation of SESN2 in 293T cells. **f**) Western blot analysis of time-dependent degradation of SESN2 in MDA-MB-231 and MDA-MB-468 cells.

In addition, we also found some CC-885-induced CRBN-interactors were not degraded substrates (Supplementary table 6), which is also observed in previous studies.^50, 52^ This is reasonable since there are lots of factors that could influence the cellular degradation kinetics and process,^53.^ such as the dynamic polyubiquitylation and correct recognition by proteasome, etc. Of note, such PPI information is still important. First, these interactions might provide insights into how molecular glue mediates protein recognitions when these protein complexes are resolved by structural biology. In addition, since the role of CRBN is to recruit substrates for ubiquitylation by CRL4^CRBN^ E3 ligase, the non-degradation substrates of CRBN may still undergo ubiquitylation, which might lead to functional alteration of the target proteins and thus diverse physiological effects.^54^ Interestingly, other substrates such as IKZF1/3 previously found in multiple myeloma cell lines are not enriched, mainly due to the low protein abundance of these proteins in 293T cell line (according to the Human Protein Atlas database),^55^ which reflects that the distinct substrate profile could be obtained in different cell lines.^56^ Altogether, by applying our VL-XL strategy, we not only successfully retrieved known substrates of MG CC-885 but also identified its new degradation substrate, suggesting the power of our method in deciphering MG-induced dynamic PPIs and substrate profile of MG degrader in live cells. Thus, our strategy would open an innovation avenue for the design of new MGs to modulate/redirect the function of the substrates of the MG-induced PPIs.

### Advancements of VL-XL strategy for protein interaction studies

We have demonstrated the VL-CO cross-linker is superior for capturing and enabling in-depth analysis of PPIs, and the visible-light-driven cross-linking strategy is priority choice for cross-linking and analysis of PPIs *in vitro* and in living cells. In our study profiling the EGF-stimulated dynamic EGFR interactome, and the substrate profile of the E3 ligase CRBN induced by the MG degrader, we employed a visible-light-induced cross-linking strategy to capture dynamic PPIs, thereby facilitating a precise analysis of protein interactions in native biological environments.

In fact, although PANAC photoclick chemistry has been employed in a variety of applications across several fields,^24-37^ these applications have so far been exclusively triggered by UV light (Figure 1b). Moreover, bifunctional photo-cross-linkers for XL-MS studies based on PANAC photoclick chemistry have not yet been developed. In this study, leveraging the mechanism of PANAC photoclick chemistry, we have successfully developed visible-light-responsive bifunctional photo-cross-linkers for XL-MS studies (Figure 1c and 2). Our visible-light-responsive photo-cross-linkers and the VL-XL strategy differ significantly from previous applications of PANAC photoclick chemistry in terms of reactive functionality, light source, chemical structure, and application (Figure 3, 4, 5).^24-37^ Our approach enables visible-light-induced lysine-specific cross-linking for MS analysis, representing an innovative photo-cross-linking strategy in the XL-MS field. More importantly, the use of visible light offers superior biocompatibility, and yet, visible-light-induced PANAC photoclick chemistry has remained underdeveloped and is highly desired. In this context, our work introduces visible-light-responsive reactive functionalities (Figure 2) and may inspire innovative solutions for developing the next generation of PANAC photoclick chemistry through visible-light activation. Consequently, our research paves the way for future advancements in visible-light-induced PANAC photoclick chemistry, offering exciting opportunities for diverse applications across various fields.

## Conclusion

In summary, we have developed VL-XL strategy for efficient cross-linking of protein complexes both *in vitro* and in living cells, enabling in-depth analysis of protein structures and dynamic PPIs. By harnessing the advantages of temporal controllability, lysine-selectivity and good biocompatibility, our VL-XL strategy is highly accessible for MS data analysis and simplifies the cross-linking sites mapping, thus facilitates obtaining accurate information of protein structures and deciphering elusive PPIs in living systems. Building on these significant advancements, we demonstrate the capability of the VL-XL strategy to investigate the dynamic PD-L1 dimerization stimulated by an exogenous modulator, and profile time-resolved EGF-stimulated dynamic EGFR interactome providing valuable insights into the regulatory mechanisms of EGFR signaling and intracellular trafficking. Moreover, the VL-XL strategy effectively deciphers MG-induced dynamic PPIs and substrate profile of MG degrader in live cells, opening an innovative avenue for identifying neo-substrates of MG degraders.

Collectively, our VL-XL strategy expands the XL-MS toolbox by providing a reliable cross-linking method for precise analysis of protein complexes and in-depth investigation of dynamic PPIs in live cells. This approach addresses the challenges associated with lacking temporal controllability, UV-dependent and non-selective cross-linking manners, which will inspire effective solutions for XL-MS studies. We anticipate that future efforts to incorporate cleavable or enrichable spacers into visible-light-responsive cross-linkers will lead to broad applications, facilitating the exploration of ambitious biological questions in fields of protein interactomes, chemical biology and structural systems biology.

## Supporting Information

Supplementary figures (S1-21). ^1^H and ^13^C NMR spectra for all compounds. Additional experimental details, materials and methods. Supplementary table.

## Notes

The authors declare no competing financial interest.

## Acknowledgements

This study was supported by the National Science Foundation of China (No. 22307126, 22377136, 92053106, 22225702, 22377133, T2488301).

## References

(1) Chavez, J. D.; Wippel, H. H.; Tang, X.; Keller, A.; Bruce, J. E. In-Cell Labeling and Mass Spectrometry for Systems-Level Structural Biology. Chem. Rev. 2022, 122 (8), 7647–7689.

(2) O’Reilly, F. J.; Rappsilber, J. Cross-linking mass spectrometry: methods and applications in structural, molecular and systems biology. Nat. Struct. Mol. Biol. 2018, 25 (11), 1000–1008.

(3) Kustatscher, G.; Collins, T.; Gingras, A. C.; Guo, T.; Hermjakob, H.; Ideker, T.; Lilley, K. S.; Lundberg, E.; Marcotte, E. M.; Ralser, M.; et al. Understudied proteins: opportunities and challenges for functional proteomics. Nat. Methods 2022, 19 (7), 774–779.

(4) Yu, C.; Huang, L. Cross-Linking Mass Spectrometry: An Emerging Technology for Interactomics and Structural Biology. Anal. Chem. 2018, 90 (1), 144–165.

(5) Piersimoni, L.; Kastritis, P. L.; Arlt, C.; Sinz, A. Cross-Linking Mass Spectrometry for Investigating Protein Conformations and Protein-Protein Interactions─A Method for All Seasons. Chem. Rev. 2022, 122 (8), 7500–7531.

(6) Iacobucci, C.; Piotrowski, C.; Aebersold, R.; Amaral, B. C.; Andrews, P.; Bernfur, K.; Borchers, C.; Brodie, N. I.; Bruce, J. E.; Cao, Y.; et al. First Community-Wide, Comparative Cross-Linking Mass Spectrometry Study. Anal. Chem. 2019, 91 (11), 6953–6961.

(7) Jin, Z.; Wan, L.; Zhang, Y.; Li, X.; Cao, Y.; Liu, H.; Fan, S.; Cao, D.; Wang, Z.; Li, X.; et al. Structure of a TOC-TIC supercomplex spanning two chloroplast envelope membranes. Cell 2022, 185 (25), 4788–4800.e4713.

(8) Guan, H.; Wang, P.; Zhang, P.; Ruan, C.; Ou, Y.; Peng, B.; Zheng, X.; Lei, J.; Li, B.; Yan, C.; et al. Diverse modes of H3K36me3-guided nucleosomal deacetylation by Rpd3S. Nature 2023, 620 (7974), 669–675.

(9) Gonzalez-Lozano, M. A.; Koopmans, F.; Sullivan, P. F.; Protze, J.; Krause, G.; Verhage, M.; Li, K. W.; Liu, F.; Smit, A. B. Stitching the synapse: Cross-linking mass spectrometry into resolving synaptic protein interactions. Sci. Adv. 2020, 6 (8), eaax5783.

(10) Schweppe, D. K.; Chavez, J. D.; Lee, C. F.; Caudal, A.; Kruse, S. E.; Stuppard, R.; Marcinek, D. J.; Shadel, G. S.; Tian, R.; Bruce, J. E. Mitochondrial protein interactome elucidated by chemical cross-linking mass spectrometry. Proc. Natl. Acad. Sci. U.S.A. 2017, 114 (7), 1732–1737.

(11) Wheat, A.; Yu, C.; Wang, X.; Burke, A. M.; Chemmama, I. E.; Kaake, R. M.; Baker, P.; Rychnovsky, S. D.; Yang, J.; Huang, L. Protein interaction landscapes revealed by advanced in vivo cross-linking-mass spectrometry. Proc. Natl. Acad. Sci. U.S.A. 2021, 118 (32), e2023360118.

(12) Chen, J.; Zhao, Q.; Gao, H.; Zhao, L.; Chu, H.; Shan, Y.; Liang, Z.; Zhang, Y.; Zhang, L. A Glycosidic-Bond-Based Mass-Spectrometry-Cleavable Cross-linker Enables In Vivo Cross-linking for Protein Complex Analysis. Angew. Chem. Int. Ed. 2023, 62 (24), e202212860.

(13) Jiang, P.-L.; Wang, C.; Diehl, A.; Viner, R.; Etienne, C.; Nandhikonda, P.; Foster, L.; Bomgarden, R. D.; Liu, F. A Membrane-Permeable and Immobilized Metal Affinity Chromatography (IMAC) Enrichable Cross-Linking Reagent to Advance In Vivo Cross-Linking Mass Spectrometry. Angew. Chem. Int. Ed. 2022, 61 (12), e202113937.

(14) Leitner, A.; Walzthoeni, T.; Kahraman, A.; Herzog, F.; Rinner, O.; Beck, M.; Aebersold, R. Probing Native Protein Structures by Chemical Cross-linking, Mass Spectrometry, and Bioinformatics. Mol. Cell. Proteomics 2010, 9 (8), 1634–1649.

(15) Mädler, S.; Bich, C.; Touboul, D.; Zenobi, R. Chemical cross-linking with NHS esters: a systematic study on amino acid reactivities. J. Mass Spectrom. 2009, 44 (5), 694–706.

(16) Kumar, G. S.; Lin, Q. Light-Triggered Click Chemistry. Chem. Rev. 2021, 121, 6991–7031.

(17) Preston, G. W.; Wilson, A. J. Photo-induced covalent cross-linking for the analysis of biomolecular interactions. Chem. Soc. Rev. 2013, 42 (8), 3289–3301.

(18) Coin, I. Application of non-canonical crosslinking amino acids to study protein–protein interactions in live cells. Curr. Opin. Chem. Biol. 2018, 46, 156–163.

(19) Liu, J.; Cai, L.; Sun, W.; Cheng, R.; Wang, N.; Jin, L.; Rozovsky, S.; Seiple, I. B.; Wang, L. Photocaged Quinone Methide Crosslinkers for Light-Controlled Chemical Crosslinking of Protein–Protein and Protein–DNA Complexes. Angew. Chem. Int. Ed. 2019, 58 (52), 18839–18843.

(20) Pehrson, J. R. Thymine dimer formation as a probe of the path of DNA in and between nucleosomes in intact chromatin. Proc. Natl. Acad. Sci. U.S.A. 1989, 86 (23), 9149–9153.

(21) Pattison, D. I.; Rahmanto, A. S.; Davies, M. J. Photo-oxidation of proteins. Photochem. Photobiol. Sci. 2012, 11 (1), 38–53.

(22) Sinha, N. K.; McKenney, C.; Yeow, Z. Y.; Li, J. J.; Nam, K. H.; Yaron-Barir, T. M.; Johnson, J. L.; Huntsman, E. M.; Cantley, L. C.; Ordureau, A.; et al. The ribotoxic stress response drives UV-mediated cell death. Cell 2024, 187 (14), 3652–3670.

(23) Ryu, K. A.; Kaszuba, C. M.; Bissonnette, N. B.; Oslund, R. C.; Fadeyi, O. O. Interrogating biological systems using visible-light-powered catalysis. Nat. Rev. Chem. 2021, 5 (5), 322–337.

(24) Guo, A.-D.; Wei, D.; Nie, H.-J.; Hu, H.; Peng, C.; Li, S.-T.; Yan, K.-N.; Zhou, B.-S.; Feng, L.; Fang, C.; et al. Light-induced primary amines and o-nitrobenzyl alcohols cyclization as a versatile photoclick reaction for modular conjugation. Nat. Commun. 2020, 11 (1), 5472.

(25) Hu, W.; Yuan, Y.; Wang, C.-H.; Tian, H.-T.; Guo, A.-D.; Nie, H.-J.; Hu, H.; Tan, M.; Tang, Z.; Chen, X.-H. Genetically Encoded Residue-Selective Photo-Crosslinker to Capture Protein-Protein Interactions in Living Cells. Chem 2019, 5 (11), 2955–2968.

(26) Guo, A.-D.; Yan, K.-N.; Hu, H.; Zhai, L.; Hu, T.-F.; Su, H.; Chi, Y.; Zha, J.; Xu, Y.; Zhao, D.; et al. Spatiotemporal and global profiling of DNA–protein interactions enables discovery of low-affinity transcription factors. Nat. Chem. 2023, 15 (6), 803–814.

(27) Hu, H.; Hu, W.; Guo, A.-D.; Zhai, L.; Ma, S.; Nie, H.-J.; Zhou, B.-S.; Liu, T.; Jia, X.; Liu, X.; et al. Spatiotemporal and direct capturing global substrates of lysine-modifying enzymes in living cells. Nat. Commun. 2024, 15 (1), 1465.

(28) Zanon, P. R. A.; Yu, F.; Musacchio, P.; Lewald, L.; Zollo, M.; Krauskopf, K.; Mrdović, D.; Raunft, P.; Maher, T. E.; Cigler, M.; et al. Profiling the proteome-wide selectivity of diverse electrophiles. ChemRxiv [Preprint] 2021, doi:10.26434/chemrxiv-2021-w7rss-v2.

(29) Nie, H.-J.; Guo, A.-D.; Lin, H.-X.; Chen, X.-H. Rapid and halide compatible synthesis of 2-N-substituted indazolone derivatives via photochemical cyclization in aqueous media. RSC Adv. 2019, 9 (23), 13249–13253.

(30) Yan, K.-N.; Nie, Y.-Q.; Wang, J.-Y.; Yin, G.-L.; Liu, Q.; Hu, H.; Sun, X.; Chen, X.-H. Accelerating PROTACs Discovery Through a Direct-to-Biology Platform Enabled by Modular Photoclick Chemistry. Adv. Sci. 2024, 11 (26), 2400594.

(31) Li, J.; Hu, Q.-L.; Song, Z.; Chan, A. S. C.; Xiong, X.-F. Cleavable Cys labeling directed Lys site-selective stapling and single-site modification. Sci. China Chem. 2022, 65 (7), 1356–1361.

(32) Guo, A.-D.; Wu, K.-H.; Chen, X.-H. Light-induced efficient and residue-selective bioconjugation of native proteins via indazolone formation. RSC Adv. 2021, 11 (4), 2235–2241.

(33) Suo, Y.; Li, K.; Ling, X.; Yan, K.; Lu, W.; Yue, J.; Chen, X.; Duan, Z.; Lu, X. Discovery Small-Molecule p300 Inhibitors Derived from a Newly Developed Indazolone-Focused DNA-Encoded Library. Bioconj. Chem. 2024, 35 (8), 1251–1257.

(34) Wei, H.; Zhang, T.; Li, Y.; Zhang, G.; Li, Y. Covalent Capture and Selection of DNA-Encoded Chemical Libraries via Photo-Activated Lysine-Selective Crosslinkers. Chem. Asian J. 2023, 18 (21), e202300652.

(35) Lu, D.-Q.; Liu, D.; Liu, J.; Li, W.-X.; Ai, Y.; Wang, J.; Guan, D. Facile synthesis of chitosan-based nanogels through photo-crosslinking for doxorubicin delivery. Int. J. Biol. Macromol. 2022, 218, 335–345.

(36) Li, M.; Liu, X.; Shang, J.; Wang, X.; Zhang, X.-B.; Xiong, B. Light-mediated protein functionalization of photoclickable hydrogel interface for selective cell capture and dot blotting assay. Talanta 2024, 267, 125248.

(37) Wang, J.; Qi, F.; Feng, H.; Xu, A.; Lu, D.-Q.; Liang, J.; Zhang, Z.; Li, J.; Liu, D.; Zhang, B.; et al. In situ formed tissue-adhesive carboxymethyl chitosan hydrogels through photoclick chemistry for wound healing. Eur. Polym. J. 2024, 203, 112680.

(38) Zhu, C.; Kou, T.; Kadi, A. A.; Li, J.; Zhang, Y. Molecular platforms based on biocompatible photoreactions for photomodulation of biological targets. Org. Biomol. Chem. 2021, 19 (43), 9358–9368.

(39) Tomas, A.; Futter, C. E.; Eden, E. R. EGF receptor trafficking: consequences for signaling and cancer. Trends Cell Biol. 2014, 24 (1), 26–34.

(40) Chen, Y.; Leng, M.; Gao, Y.; Zhan, D.; Choi, J. M.; Song, L.; Li, K.; Xia, X.; Zhang, C.; Liu, M.; et al. A Cross-Linking-Aided Immunoprecipitation/Mass Spectrometry Workflow Reveals Extensive Intracellular Trafficking in Time-Resolved, Signal-Dependent Epidermal Growth Factor Receptor Proteome. J. Proteome Res. 2019, 18 (10), 3715–3730.

(41) Goley, E. D.; Welch, M. D. The ARP2/3 complex: an actin nucleator comes of age. Nat. Rev. Mol. Cell Biol. 2006, 7 (10), 713–726.

(42) Liu, J.-J. Regulation of dynein-dynactin-driven vesicular transport. Traffic 2017, 18 (6), 336–347.

(43) Márquez, M. G.; Brandán, Y. R.; Guaytima, E. d. V.; Paván, C. H.; Favale, N. O.; Sterin-Speziale, N. B. Physiologically induced restructuring of focal adhesions causes mobilization of vinculin by a vesicular endocytic recycling pathway. Biochim. Biophys. Acta 2014, 1843 (12), 2991–3003.

(44) Abramson, J., Adler, J. Dunger, J. Evans, R. Green, T. Pritzel, A. Ronneberger, O. Willmore, L. Ballard, A Bambrick, J. et al. Accurate structure prediction of biomolecular interactions with AlphaFold 3. Nature 2024, 630, 493–500.

(45) Schreiber, S. L. The Rise of Molecular Glues. Cell 2021, 184 (1), 3–9.

(46) Kozicka, Z.; Thomä, N. H. Haven’t got a glue: Protein surface variation for the design of molecular glue degraders. Cell Chem. Biol. 2021, 28 (7), 1032–1047.

(47) Duran-Frigola, M.; Cigler, M.; Winter, G. E. Advancing Targeted Protein Degradation via Multiomics Profiling and Artificial Intelligence. J. Am. Chem. Soc. 2023, 145 (5), 2711–2732.

(48) Dong, G.; Ding, Y.; He, S.; Sheng, C. Molecular Glues for Targeted Protein Degradation: From Serendipity to Rational Discovery. J. Med. Chem. 2021, 64 (15), 10606–10620.

(49) Matyskiela, M. E.; Lu, G.; Ito, T.; Pagarigan, B.; Lu, C.-C.; Miller, K.; Fang, W.; Wang, N.-Y.; et al. A novel cereblon modulator recruits GSPT1 to the CRL4CRBN ubiquitin ligase. Nature 2016, 535 (7611), 252–257.

(50) Huang, H.-T.; Lumpkin, R. J.; Tsai, R. W.; Su, S.; Zhao, X.; Xiong, Y.; Chen, J.; Mageed, N.; Donovan, K. A.; Fischer, E. S.; et al. Ubiquitin-specific proximity labeling for the identification of E3 ligase substrates. Nat. Chem. Biol. 2024, 20 (9), 1227–1236.

(51) Chen, Y.; Huang, T.; Yu, Z.; Yu, Q.; Wang, Y.; Hu, J. a.; Shi, J.; Yang, G. The functions and roles of sestrins in regulating human diseases. Cell. Mol. Biol. Lett. 2022, 27 (1), 2.

(52) Sievers, Q. L.; Petzold, G.; Bunker, R. D.; Renneville, A.; Słabicki, M.; Liddicoat, B. J.; Abdulrahman, W.; Mikkelsen, T.; Ebert, B. L.; Thomä, N. H. Defining the human C2H2 zinc finger degrome targeted by thalidomide analogs through CRBN. Science 2018, 362 (6414), eaat0572.

(53) Riching, K. M.; Caine, E. A.; Urh, M.; Daniels, D. L. The importance of cellular degradation kinetics for understanding mechanisms in targeted protein degradation. Chem. Soc. Rev. 2022, 51 (14), 6210–6221.

(54) Damgaard, R. B. The ubiquitin system: from cell signalling to disease biology and new therapeutic opportunities. Cell Death Differ. 2021, 28 (2), 423–426.

(55) Uhlen, M.; Oksvold, P.; Fagerberg, L.; Lundberg, E.; Jonasson, K.; Forsberg, M.; Zwahlen, M.; Kampf, C.; Wester, K.; Hober, S.; et al. Towards a knowledge-based Human Protein Atlas. Nat. Biotechnol. 2010, 28 (12), 1248–1250.

(56) Oleinikovas, V.; Gainza, P.; Ryckmans, T.; Fasching, B.; Thomä, N. H. From Thalidomide to Rational Molecular Glue Design for Targeted Protein Degradation. Annu. Rev. Pharmacol. Toxicol. 2024, 64, 291–312.

